# The importance of being cross-linked for the bacterial cell wall

**DOI:** 10.1101/573113

**Authors:** Garima Rani, Issan Patri

**Affiliations:** The Institute of Mathematical Sciences, CIT Campus, Taramani, Chennai, India; Homi Bhabha National Institute, Training School Complex, Anushaktinagar, Mumbai 400094, India; Chennai Mathematical Institute, SIPCOT IT Park, Siruseri, Chennai, India

## Abstract

The bacterial cell wall is primarily composed of a mesh of stiff glycan strands cross-linked by peptide bridges and is essential for safeguarding the cell. The structure of the cell wall has to be stiff enough to bear the high turgor pressure and sufficiently tough to ensure protection against failure. Here we explore the role of various design features of the cell in enhancing the toughness of the cell wall. We explain how the glycan strand length distribution and the degree of cross-linking can play a vital role in ensuring that the cell wall offers sufficient resistance to propagation of cracks. We suggest a possible mechanism by which peptide bond hydrolysis can also help mitigate this risk of failure. We also study the reinforcing effect of MreB on the cell wall and conclude that the cross-linked structure of the cell wall plays the more important role in safeguarding against mechanical failure due to cracking.

## 1 Introduction

Understanding the design features of bacterial cells has long been a fundamental topic of research and a key design feature of bacteria is the cell wall. The bacterial cell wall is primarily composed of a sturdy mesh of stiff glycan strands, crosslinked by short peptide bonds, known as the peptidoglycan network. The cell wall plays multiple roles, including protecting the cell against external threats and providing the cell its characteristic shape [1, 2, 3, 4]. It has traditionally been an important target of antibiotics like penicillin and its derivatives. However, as several strains of bacteria are becoming resistant to antibiotics [5, 6], understanding the structure of the bacterial cell wall assumes renewed significance, so that newer anti-bacterial agents that target the cell wall can be designed.

Toughness, or resistance to propagation of cracks, is an immensely desirable materials property [7]. A common problem encountered when materials are engineered is to ensure not only strength, specifically stiffness, but also toughness, which can be difficult as these two requirements are often at odds [8]. Biological materials in particular have to be structurally strong enough to resist high tension forces and sufficiently tough to prevent failure due to cracking. Indeed, several biological materials like wood, bones and nacre, serve as some of the best examples satisfying this requirement, with their specific design principles being well studied in this context [9, 10, 11]. For the bacterial cell wall, it is imperative to be stiff enough to bear the high turgor pressure and maintain shape as well as being adequately tough. While the stiffness of the bacterial cell wall has been well studied [12, 13, 14, 15], our aim here is to understand the toughness of the cell wall. The cell wall, which is under high turgor pressure, can have cracks and holes on it for a variety of reasons, like for passage of nutrients. In fact, permeability of cell walls of bacteria has been much studied [16, 17, 18] and pores of size as large as 10 nm in diameter have also been observed [19]. On the other hand, in bacteria like *Staphylococcus aureus*, mechanical crack propagation has been shown to drive daughter cell separation [20], which indicates that bacterial cells are adapted to be able to tune mechanical failure modes. However, the relation of the fine structure of the cell wall to its toughness remains to be elucidated.

In this paper, we study the role of various design components of the cell in securing the cell wall by ensuring sufficient resistance to propagation of cracks. In particular, we examine the role of the geometry of the cell, the cross-linked structure of the cell wall and the role of the MreB cytoskeleton [21, 22] in ensuring stabilization of the cell wall against crack propagation. Our model, specifically, studies the Gram-negative rod-shaped bacteria (e.g. *Escherichia coli*), with a single layer of the peptidoglycan mesh. In short time scales relevant to the problem, the behavior of the cell wall is perfectly elastic [23, 24]. Modelling the cell both as an elastic plate and cylindrical shell, we estimate the critical crack lengths under stress due to turgor pressure. We show that cross-linking is crucial for maintaining the integrity of the cell wall, since the minimum energy needed for crack propagation, called the tearing energy, is largely controlled by the degree of cross-linking. We exhibit the important role that appropriately cross-linked shorter length glycan strands can play in enhancing the tearing energy, which gives a possible explanation of the observations on the extensive presence of short length glycan strands in the peptidoglycan mesh [25] and the strong propensity of the glycan strands to be cross-linked at the terminating units [26, 27]. Our analysis suggests that peptide bond hydrolysis can be used by the cell as a defence mechanism against cracking, since such hydrolysis can act to increase the tearing energy. Our model also provides an explanation for the observation of preferential rupturing in the circumferential direction in sonication experiments [28]. Finally, we probe the role of the MreB cytoskeleton which can promote cell wall toughness by exerting an inward directed pressure countering the turgor pressure. Our results indicate that the cross-linked structure of the cell wall plays the more important role in safeguarding the cell against failure due to crack propagation.

## 2 Cell Wall Model

We model the cell wall as a pressurized, linear elastic, thin cylindrical shell, as depicted in Figure 1. We assume for simplicity that the shell is isotropic with elastic modulus denoted *E*. One can consider an orthotropic cell wall model [29] by modifying the strain energy release term (see Section 3) for which additional elastic constants are required, which is discussed in Appendix B. The radius of the shell is *R* and its thickness, *h*. Since *h/R* is small (relevant parameter values are listed in Table 2), we can treat the cell wall as effectively two dimensional, describing it by its central (neutral) surface. We use the thin shell “membrane” approximation, neglecting all moment expressions [30]. The turgor pressure is denoted *P*. Using force-balance, the hoop stress is

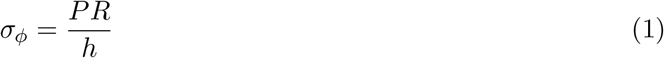

while the axial stress is

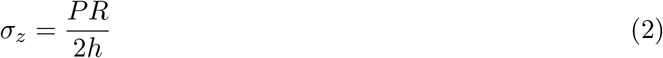

Hence a stress anisotropy exists in the cylindrical case, with *σ*_*φ*_/*σ*_*z*_ = 2.

**Figure 1:**
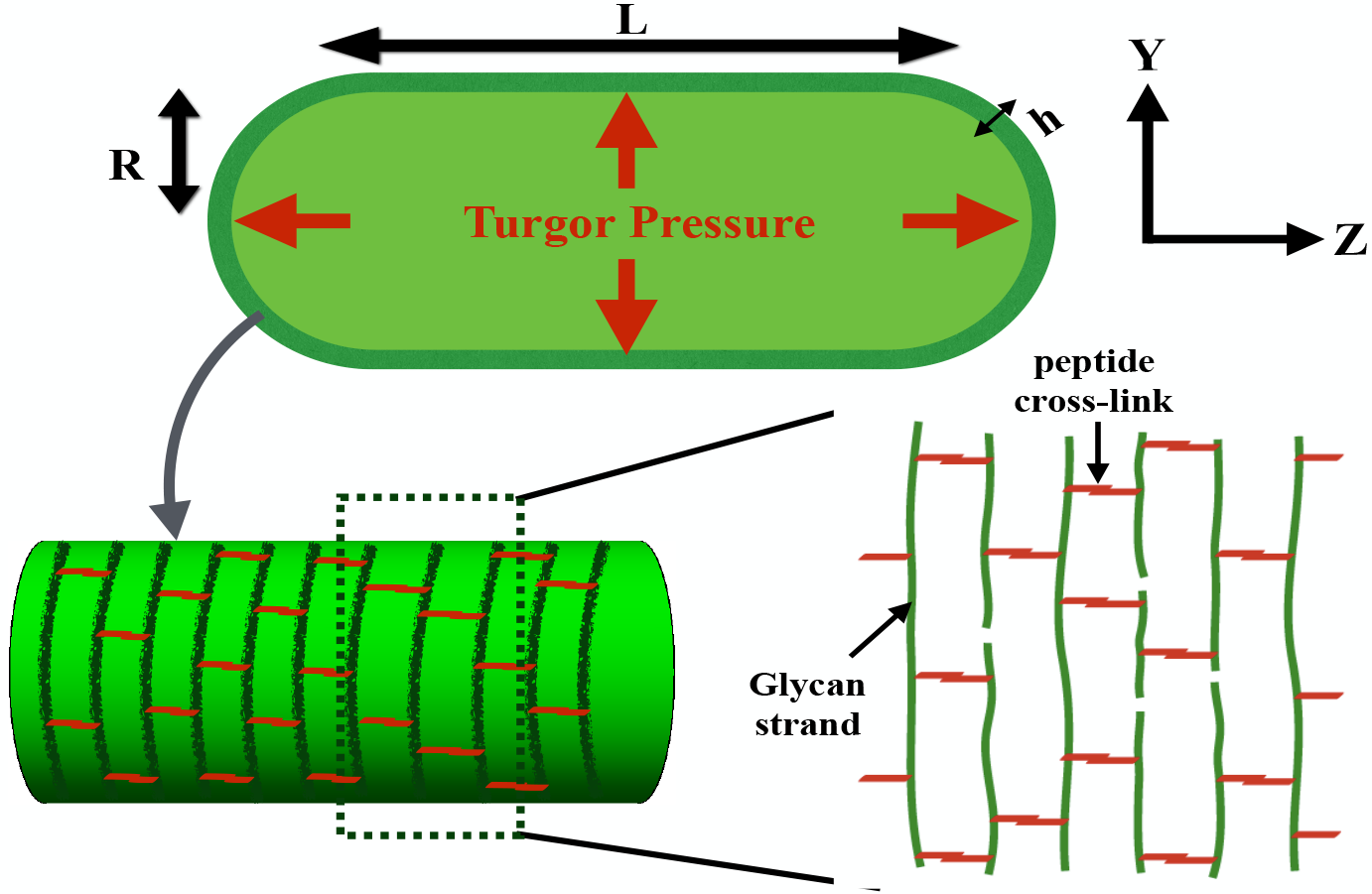
The bacterial cell wall modelled as a pressurised cylindrical shell of radius *R*, thickness *h* and with the length denoted *L*. Stiff glycan strands in the cell wall, aligned roughly in the circumferential direction, are cross-linked by peptide bonds.

## 3 Cell Wall Crack Energetics

Since stress in the hoop direction is twice the stress in the axial direction, longitudinal cracks are subject to a larger stress, as they are aligned perpendicular to the hoop direction. Indeed, cylindrical pressure vessels predominantly display longitudinal cracks, for this reason, including well known examples occurring in daily life like sausages and pipes cracking longitudinally [31]. For rod shaped bacteria, like *E.coli*, which is under high turgor pressure, a corresponding stress anisotropy indicates the possibility of similar failure due to cracking in the longitudinal direction. Our aim here is to understand the structural features of the cell wall which protect it from such failure. We model a crack on the cell wall in two ways: (1) by considering a centrally placed crack on an infinite plate, thus neglecting curvature effects, and (2) considering a longitudinally aligned crack on a pressurized shell, thus accounting for cell curvature.

In the first case, we begin with a thin plate of thickness *h* placed in the *Y Z*-plane, with a tensile load *σ* = *PR/h* applied in the *Y* -direction. A crack of length 2*c* is introduced, along the *Z*-axis, as shown in Fig. 2(a). The crack lengths of interest to us are considerably smaller than the radius and length of the cell wall. We can thus treat the plate as infinite. To calculate the critical crack length, beyond which crack propagation becomes energetically favourable, we use the Griffith’s criterion. The Griffith’s criterion [32] compares the energy required to break atomic bonds, thus leading to new surfaces, to the strain energy released as the crack enlarges (assuming that no energy dissipation occurs). The strain energy released in this case is given by

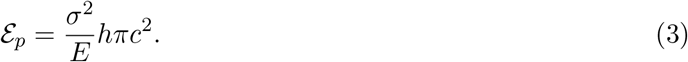

**Figure 2:**
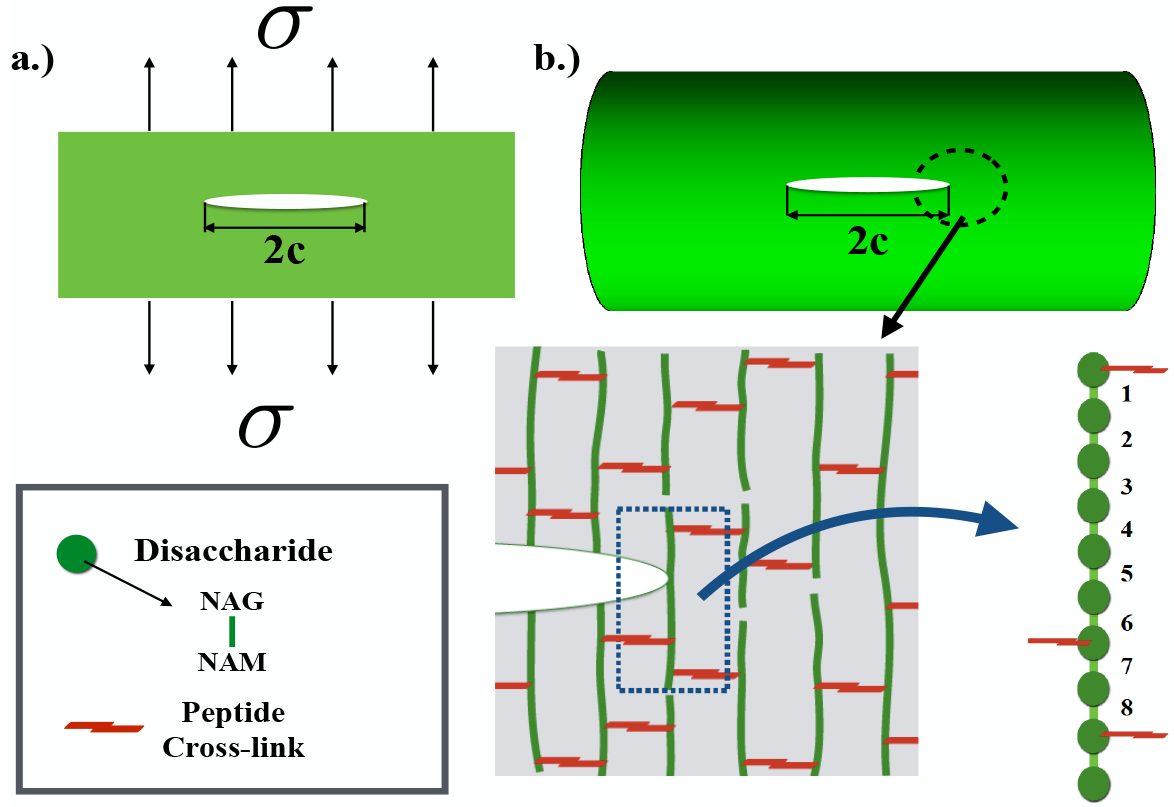
(a) A crack of length 2*c* on a thin plate of thickness *h* with a remote load *σ* acting on it, perpendicular to the line of crack. (b) A longitudinally aligned crack on the cell wall. Near the tip, the crack comes up against a glycan strand, here illustrated having a length of 10 disaccharide units, with average number of glycosidic bonds connecting the disaccharides between adjacent cross-links calculated as *n* = 4. Each disaccharide unit consists of alternating sugars NAG and NAM connected by a glycosidic bond.

In the case of an orthotropic model of the cell wall, this term has to be modified with additional elastic constants incorporated into the expression (in Appendix B, we show that under reasonable assumptions, the strain energy released in the anisotropic model is comparable to the isotropic model used here).

The critical crack length is given by

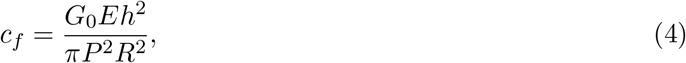

where *G*_0_ denotes the minimum tearing energy (see Appendix A). We note that this paradigm of modelling curved objects by flat surfaces has been utilized widely, both in the case of bacteria [33] and other cells [34].

In the second case, we incorporate the cell curvature into our calculation. We consider a longitudinally aligned crack of length 2*c* on a thin, pressurized, cylindrical shell a shown in (Figure 2(b)). In this case, the effect of geometry and the internal pressure results in an out-of-plane deformation of the shell in the periphery of the crack, due to which an additional strain energy is released, as compared to crack on a plate [35, 36, 37]. In Appendix C, we give a simple derivation of the total strain energy released in this case as

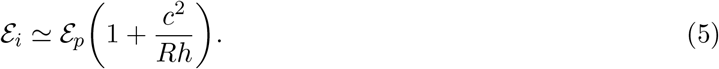

We can now calculate the critical crack length *c*_*f*_ using the Griffith’s criterion, as the unique (positive) real root of the cubic equation

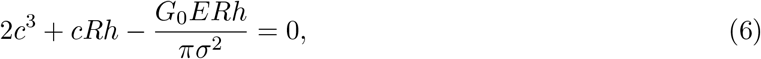

where *σ* = *σ*_*φ*_ = *PR*/*h*.

To understand the effect of geometry of the cell wall, a comparison of these two cases needs to made. For this, we first need to compute the minimum tearing energy *G*_0_, which we do in the following section.

### 3.1 Tearing Energy: Role of Cross-Links

A crack typically propagates by rupturing the bonds lying across its plane [38]. Thus, the tearing energy is conventionally calculated by estimating the energy cost for disrupting the bonds lying across the plane of the crack. However, in a cross-linked polymer, as the crack propagates, it encounters chains of monomers lying between adjacent cross-links. The forces involved are transmitted through the cross-links. So, in order to break the chain, each bond in the chain has to be supplied energy almost equalling the energy required to rupture them even though one of the bonds might eventually rupture [39, 40]. Such cross-linking thus acts as a bulwark for the polymer, acting to protect it from mechanical failure due to cracking.

For an axially aligned crack on the cell wall to propagate, it must cross a number of glycan strands cross-linked by peptide bridges. Glycan strands are made up of repeating disaccharide units of N-acetylglucosamine (NAG) and N-acetylmuramic acid (NAM). A peptide stem of few amino acids is attached to NAM. The glycosidic bonds between the alternating sugars NAG and NAM form the backbone of the glycan chain [41]. So, if there are on an average *m* such bonds between adjacent cross links, then total energy needed to disrupt the chain will be ≈ *mE_c_*, where *E*_*c*_ denotes the dissociation energy of the bond. Let Σ denote the number of glycan strands crossing per unit area in the fracture plane. We then have

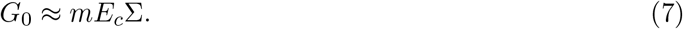

To relate *m* to the average length of the glycan chain between adjacent cross-links (denoted *l_a_*), we define *n* to be the average number of glycosidic bonds connecting the disaccharide units between adjacent cross-links (see Fig. 2(b)). It follows then that *m* = 2*n* and *l*_*a*_ = (*n* + 1) disaccharides.

We estimate *G*_0_ as follows- the average dissociation energy of a glycosidic bond, specifically, a C-O bond, is of the order of *E*_*c*_ ≈ 6 × 10^−19^*J*. The thickness of the cell wall is ≈ 5 nm and the glycan inter-strand spacing is ≈ 2 nm (see Table 2). We then get Σ ≈ (6 × 10^16^/*m*^2^). Next we estimate *n* across a given glycan strand. We have

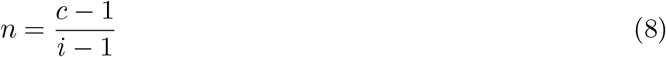

where *c* denotes the total number of disaccharide units between the two extreme cross-links in the glycan strand and *i* denotes the total number of cross-linked peptide stems across the length of the strand. As explained in Appendix D, in the limit of long glycan strands spanning the circumference of the cell, we have *n* ~ 1/*k*, where *k* denotes the degree of cross-linking across the glycan strand. In case of *E.coli*, peptide stems rotate around the glycan backbone by roughly 90° per disaccharide [4], so only 50% of the peptide stems are available for cross-linking and it has been observed that the degree of cross-linking is around 30% [26], so *n* ∼ 3. However, as discussed in Appendix D, placing of the cross-links at the terminating units can increase the value of *n* for shorter length glycan strands substantially. In particular, for glycan strands which are cross-linked at the ends, the value of the numerator in Equation 8 is maximized. For any glycan strand satisfying this property and allowing for atleast two cross-links across the strand, we get *n* ∼ 3 - 6, with strands of length 7 - 8 disaccharides being optimal in this regard, having *n* ∼ 5 - 6, while in the limit of long glycan strands, we have *n* ∼ 3. This in particular can explain following remarkable experimental observations 1- in Ref. [25], where HPLC analysis of glycan strand length distribution detected a substantial presence of short length glycan strands, with the length distribution having a mean value of ∼ 8 disaccharides for about 70% of the strands and 2- in Ref. [26, 27, 1], which concluded that ≥ 80% of the 1,6 anhydroMur*N* Ac terminal muropeptides are cross-linked and suggested similar proportions of cross-linking at the Glc*N* ac termini as inferred from peptidoglycan labelled with galactosyl transferase. Our analysis underlines the significance of terminally cross-linked short length glycan strands.

The extent of cross-linking also plays a key role in the above analysis. A high degree of cross-linking increases the stiffness of the cell wall [12]. However, it follows from Eqn. 7 and Eqn. 8 that in this case, the tearing energy is lower, thus making the structure vulnerable to cracks. On the other hand, while a lower degree of cross-linking increases the tearing energy, it also makes the cell wall less stiff [14]. Thus, the extent of cross-linking has to be delicately balanced to ensure optimal levels of stiffness and toughness.

A key mechanism of the cell wall is that of hydrolysis or the cleavage of pre-existing peptide and glycosidic bonds in the peptidoglycan mesh. Experiments indicate that *E.coli* mutants which are deficient in hydrolysis enzymes undergo rapid lysis, proving the essential role it plays in ensuring cell viability [42]. Our analysis suggests that peptide bond cleavage due to hydrolysis can secure the cell by increasing resistance to crack propagation: For the crack to propagate, it has to stretch all glycosidic bonds between adjacent cross-links before one of them ruptures. A possible way to arrest the progress of the crack is thus to cleave a peptide bond, so that the number of bonds between adjacent cross-links increases, in other words the value of *n* increases. This is analogous to the common trick employed by mechanics to arrest the progress of a crack by drilling a small hole at the tip of the crack [43]. The timescale involved in peptide bond cleavage can however impose an upper limit on the crack speed, below which bond cleavage action to inhibit crack propagation is feasible-in one cell cycle, with time *τ* = 20 minutes, a turnover of 40% - 50% of the cell wall material takes place [4] and between two adjacent circumferential cross-section of the cell, there are *≈* 500 cross-links (with 30% cross-linking and ∼ 2*πR* disaccharides comprising all glycan strands in a cross-section). So, peptide cross-links between two adjacent cross-sections are excised at a rate of *k* ∼ 10 - 12/*min*. Since a peptide bond has to be cleaved in the time that the crack front traverses the inter-strand distance *d*, the speed is thus limited to ∼ 18 - 24*nm*/*min* or slower. We have assumed that peptide bridges are cleaved uniformly across the area between the two cross-sections, although it is possible that higher stresses at the crack tip accelerate bond cleavage. Still, for a fast moving crack, peptide bond cleavage is likely to be too slow to act. For instance, in the case of *S.aureus*, daughter cell separation, for which mechanical crack propagation has been implicated, happens at speed around 1*μm/s* [20]. For crack speed of this order, peptide hydrolysis is unlikely to be able to play a mitigating role.

As we discussed, cross-linked glycan chains in the cell wall ensure that crack propagation in the longitudinal direction is effectively resisted. A natural question then is what is the preferential direction for failure under stress. It has been observed that sonication of isolated *E.coli* sacculi selectively disrupts peptide cross-links between glycan chains and rupturing tends to occur along the hoop direction [28]. This is particularly intriguing since the bond energies of glycosidic bonds and peptide bonds are very similar [28, 4]. A plausible explanation follows from our model-for a longitudinally oriented crack on the cell wall, cross-linked glycan chains crossing the fracture plane have to be taken into account for calculating the tearing energy, as explained above. On the other hand, for a crack aligned in the hoop direction to propagate, only the short peptide cross-links lying the fracture plane have to be severed. So, the tearing energy in this case is only a fraction of the longitudinal tearing energy, which can lead to a preferential rupturing in the hoop direction. A detailed study of circumferential tears in the cell wall will be carried out in a future work.

We note here that we are neglecting energy dissipation processes as the length of the dissipative zone, which is region around the crack tip where the material is no longer linear elastic and where the bulk of the energy dissipation occurs, [44, 9] is very small ∼ 5Å, as we show in Appendix E.

### 3.2 Critical Crack Length

We now calculate critical crack lengths in the planar case as well as the cylindrical case, using Griffith criterion. In Figure 3, we plot the total energy (*ε*_*t*_), defined as the difference of the strain energy released and the surface energy (see Appendix A), against the crack length, in these two cases, varying the value of *n*. The critical crack length is obtained at the maxima of the energy curve, which we observe, increases as the value of *n* increases. For the cylindrical case, the critical crack length is smaller than the planar case, with this difference increasing with increase in value of *n*. Thus at lower crack length values, the planar case provides a good approximation to the case of a crack on the cell wall under turgor pressure. However, at higher crack lengths, curvature becomes important and the planar approximation starts to break down. In the case *n* = 1, which corresponds to a 100% cross-linked sacculi, the critical crack length is small, with *c* ≈ 6*nm*. When the effect of cross-linking is completely ignored, that is when a very thin crack propagates by disrupting the bonds lying across the fracture plane (so *m* = 1 in Equation 7), then critical crack length is even smaller. This underlines the significance of appropriate levels of cross-linking in maintaining the integrity of the cell wall.

**Figure 3:**
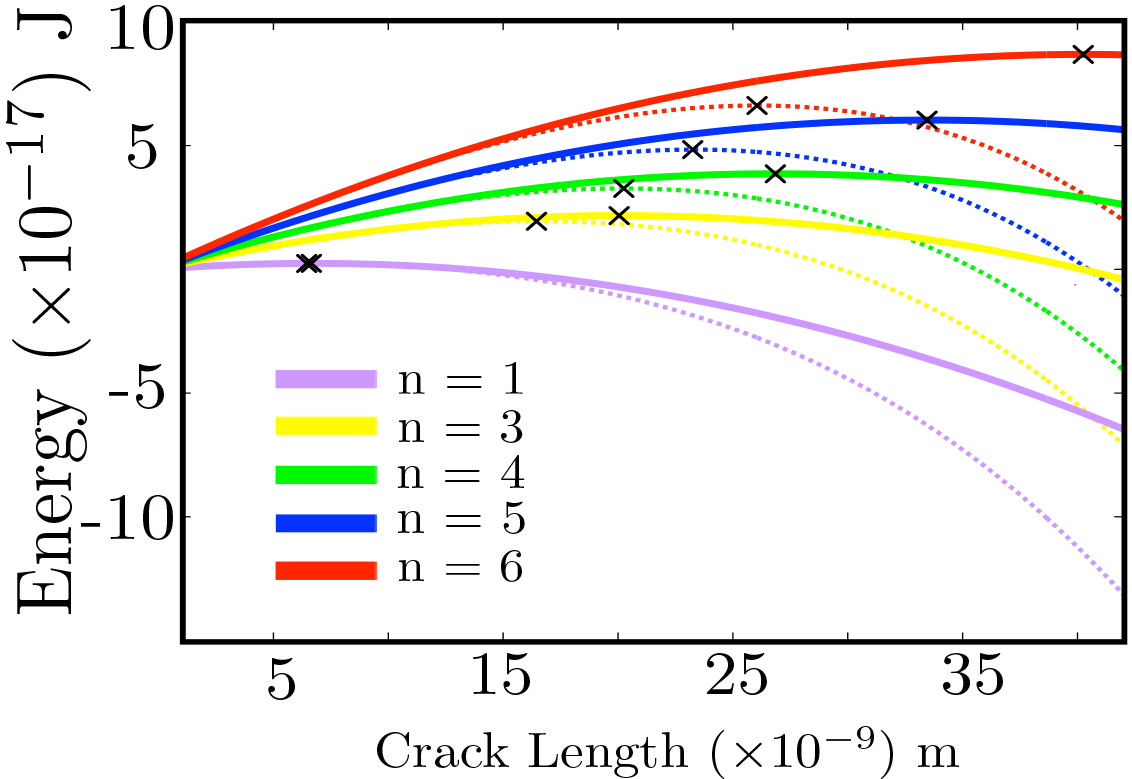
Total energy against crack length for different degrees of effect of crosslinking. The solid lines represent the plane case, while dotted lines represent the cylinder case. The cross (×) represent the critical crack lengths. For values of the elastic modulus *E*, the cylinder radius *R* and the thickness *h*, see Table 2.

Here we use Griffith’s criterion to calculate the critical crack length, which is accurately applicable for very thin cracks. For the cell wall, due to the cross-linked structure, the unstrained tip width is approximately the length between two adjacent cross-links, so the length to width ratio in our case will be ∼ 10 - 20, which ensures a degree of accuracy. More accurately, the Griffith estimate is in fact a lower bound for the critical crack length which well illustrates the reinforcing effect of the cross-linked structure of the cell wall in preventing failure due to cracking (more precise estimates of the critical crack length can be obtained by taking the exact geometry into consideration). Interestingly, critical crack length for the Gram positive bacteria *S.aureus* has been suggested to be around 40*nm* [20], which is commensurate to these estimates. Though *S.aureus* has a spherical geometry and its cell wall has a multi-layered structure, the composition of the Gram positive and Gram negative cell walls remain conserved [45], which suggests that the critical crack length in both cases can be comparable.

A natural question now is whether the role of cross-linking in maintaining the integrity of the cell can be supplanted by other design components of the cell, for instance the cytoskeleton MreB. In the next section, we explore the role of the cytoskeleton in strengthening the cell wall against failure from crack propagation. In particular, we probe if the cytoskeleton can effectively reinforce the cell against crack propagation, even when the effect of cross-linking is discounted.

## 4 Cytoskeletal Reinforcement

The actin-homologue MreB in bacteria [21, 22] is a key component of the bacterial cell. It plays an important role in the growth of the cell and in maintenance of the shape in rod shaped bacteria [46, 47]. In this section, we study the effect of MreB on the toughness of the cell wall, in particular examining the reinforcement of the cell wall by MreB and its effects on the critical crack length.

Several early papers suggested, based on *in vivo* observations, that MreB formed as a cell spanning helix [22, 48] (Figure 4 (left)). However, more recent work using high resolution light micropscopy have shown that MreB forms disconnected assemblies in the cell that move processively in the hoop direction [49, 50, 51] (Figure 4 (right)). Here, we model MreB as a collection of disconnected bent cylindrical rods oriented in the hoop direction of the cell (we consider the cell spanning helix model of MreB in Appendix H and compare it with this model).

**Figure 4:**
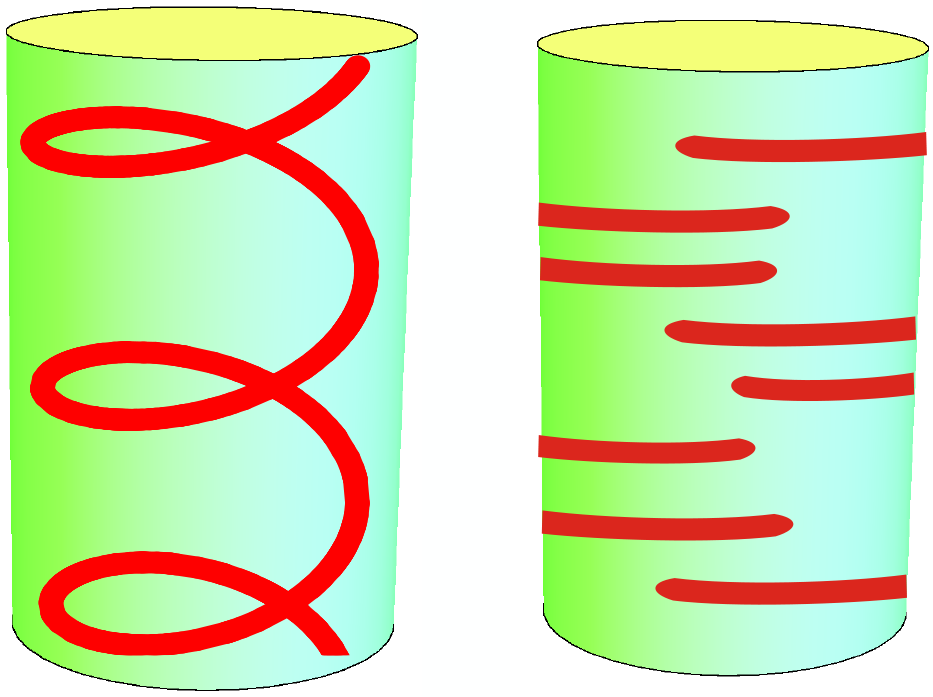
Initial studies reported actin homologue MreB modelled as a cell spanning helix (left), while newer experiments have concluded MreB forms disconnected filament assemblies aligned in the hoop direction

MreB binds directly to the cytoplasmic side of the inner membrane [52]. While the *in vivo* ultrastructure of MreB is not clear, it has been observed *in vitro* that MreB forms bundles in solutions [53] and binds as filaments to membrane [54]. Biophysical modelling has shown that the orientation of MreB along the direction of maximal curvature is determined by the trade-off between the energetics of filament bending, membrane deformation and the work done against the turgor pressure [55]. For the bacterial cell, the presence of high turgor pressure can ensure that the MreB filaments deform to the cell membrane. In this case, the configuration of MreB is distorted from its preferred shape. In vitro studies have shown that MreB filaments are highly curved (with width *>* 200 nm), so the preferred curvature is greater than the typical curvature of the bacterial cell [54, 55]. Therefore, with the preferred radius being smaller than the cell radius, an inward directed pressure is exerted, which we calculate below. It should be noted that for high values of the cross-sectional radius, the energy cost of filament bending increases and it is possible that it becomes energetically favourable for the membrane to deform to the filament [55]. In our analysis however, since we are interested in only calculating the inward pressure exerted, we take as an input a configuration of the MreB modelled as bent cylindrical rods of given cross-sectional radius, denoted *a*, which is deformed from its preferred orientation onto a given orientation determined by the cell radius. We then calculate the pressure exerted for a wide range of values of *a*, ranging from 3.2nm [54, 55] to 40nm [56]. We also note that although MreB has been observed to move persistently around the long axis of the cell over long time scales [49, 51], since we are estimating the average pressure exerted, we will assume that it is localized.

We express the energy functional as an integral over its center line, which is constrained on the cell wall. For such a curve constrained to a surface, given the Darboux frame (see Appendix F), the general energy functional reads [57]

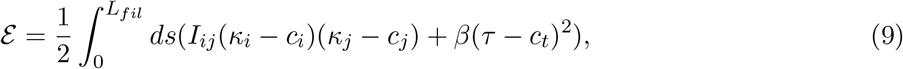

where *i, j* ∈ {*g, n*}, thus, *κ*_*i*_ and *κ*_*j*_ are from the set {*κ*_*g*_, *κ*_*n*_} of geodesic and normal curvature, and the terms *c*_*i*_, *c*_*j*_ belong to the set {*c*_*g*_, *c*_*n*_} of preferred curvatures. The tensor *I*_*ij*_ is the inertia tensor. Finally, *c*_*t*_ denotes the preferred twist and *β* denotes the twist modulus.

We assume that the cross section of MreB complex is circular. The inertia tensor *I* is thus isotropic and can be written as *I* = *αδ*_*ij*_, where 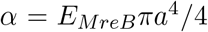 denotes the bending modulus, *a* denotes the cross-section radius and *E*_*MreB*_ the elastic modulus. We note that *β* = *α*/(1 + *ν*_*m*_) [58], where *ν*_*m*_ denotes the Poisson ratio. We make the simplifying assumption that the Poisson ratio *ν*_*m*_ = 0 for MreB, since this does not qualitatively change our results. So we have *β* = *α*.

Also, it follows from the analysis below that for a wide range of values of the radius *a*, the elastic energy is several orders of magnitude greater than *k*_*B*_*T*, hence we ignore the effect of thermal fluctuations.

### 4.1 MreB Model

MreB is modelled as several disconnected cylindrical rods, bent and oriented in the hoop direction of the cell, as shown schematically in Fig. 4(b)). We will refer to these cylindrical rods as MreB bundles. We assume that the preferred shape of MreB is a bent cylindrical rod, whose centerline can be visualised as an arc of a circle of radius *r*. In the final configuration, this radius changes to *R*. Using Eqn. 9, together with *κ*_*g*_ = *c*_*g*_ = *τ* = *c*_*t*_ = 0, *κ*_*n*_ = 1/*R* and *c*_*n*_ = 1/*r*, we obtain the total energy for *n*_*f*_ bundles as

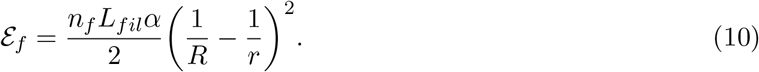

As we shall see below, the inward pressure exerted by MreB bundles is independent of their length. We thus make the simplifying assumption that the length of all the bundles is the same.

We now estimate the number of MreB bundles of a given radius and length in a typical cell. This depends on the total number of MreB monomers constituting all the MreB bundles in the cell and the number of monomers in a single bundle. We take the volume occupied by the monomers in a single bundle to be in the range (*V*_*f*_ /2, 3*V*_*f*_/4), where *V*_*f*_ denotes the volume of the bundle.

The lower bound of the packing is determined by the substantial elastic modulus of MreB (Table 2), which suggests that the monomers will have to be packed reasonably tightly, while the upper bound is determined by the well known Kepler upper bound, given by the recently proved Kepler’s conjecture, of ≈ 0.74[59]. Using this, we estimate the total number of MreB bundles in the cell (Table 1), as explained in Appendix G.

**Table 1:** Range of computed number of disconnected filament bundles in the cell for various observed values of bundle radius and lengths of filament.

| Radius \ Length | Length |  |  |
| --- | --- | --- | --- |
|  | 250 nm | 500 nm | 1500 nm |
| 3.2 nm | 437-656 | 218-328 | 73-109 |
| 10 nm | 45-67 | 22-33 | 7-11 |
| 20 nm | 11-17 | 6-9 | 2-4 |
| 40 nm | 3-5 | 1-2 | $\sim 1$ |

**Table 2:** Parameter values for *E.coli*

| Parameters | Values | References |
| --- | --- | --- |
| **Cell Wall Parameters** |  |  |
| Radius of Cell ($R$) | $\sim 0.5\mu m$ | [73] |
| Thickness of the cell wall ($h$) | $\sim 5$ nm | [74] |
| Turgor pressure ($P$) | 1 atm | [23] |
| Elastic Modulus of Cell Wall ($E$) | 30 MPa | [13] |
| Glycan Inter-strand Spacing ($d$) | 2 nm | [29] |
| **MreB Parameters** |  |  |
| Number of MreB molecules/cell ($N$) | 17000 – 40000 | [71] |
| Elastic Modulus of MreB ($E_{MreB}$) | 2 GPa (similar to actin) | [75] |
| MreB monomer diameter ($2r_0$) | 5.1 nm | [21] |

The radial force is then given by

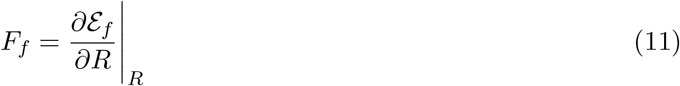

and the average pressure exerted is

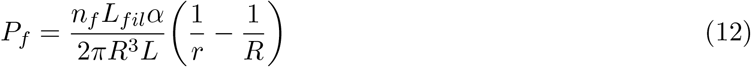

where *L* denotes the length of the cylinder. The effective turgor pressure acting on the cell wall is then given by

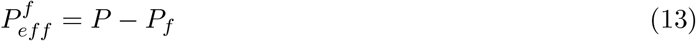

Experiments have reported shorter length MreB assemblies (∼ 250*nm*) as well as long ones nearly covering half of the cell’s circumference (∼ 1500*nm*) [55, 60, 56]. However, the pressure exerted by MreB is independent of the length of a bundle. This results from fixing the total number of MreB molecules in the cell, which effectively subsumes the role of the length of the bundles, as is derived in Appendix G. This also holds when we choose the lengths of bundles to be variable in the cell, present with varying proportions. To illustrate this, we plot the pressure exerted by the MreB against the preferred radius for different values of bundle length and fixed value of bundle radius, taken as 20 nm (left) in Figure 5. In Fig. 5 (right), we plot the pressure against preferred radius with fixed bundle length taken 250 nm and different values of bundle radius. We observe that for a wide range of preferred radius (0.2*μm* − 0.4*μm*), the pressure exerted even for very large bundle radius is much less than the turgor pressure. Similar result on the pressure exerted by helix model of MreB can also be obtained (see Appendix H, where we also compare the two models).

**Figure 5:**
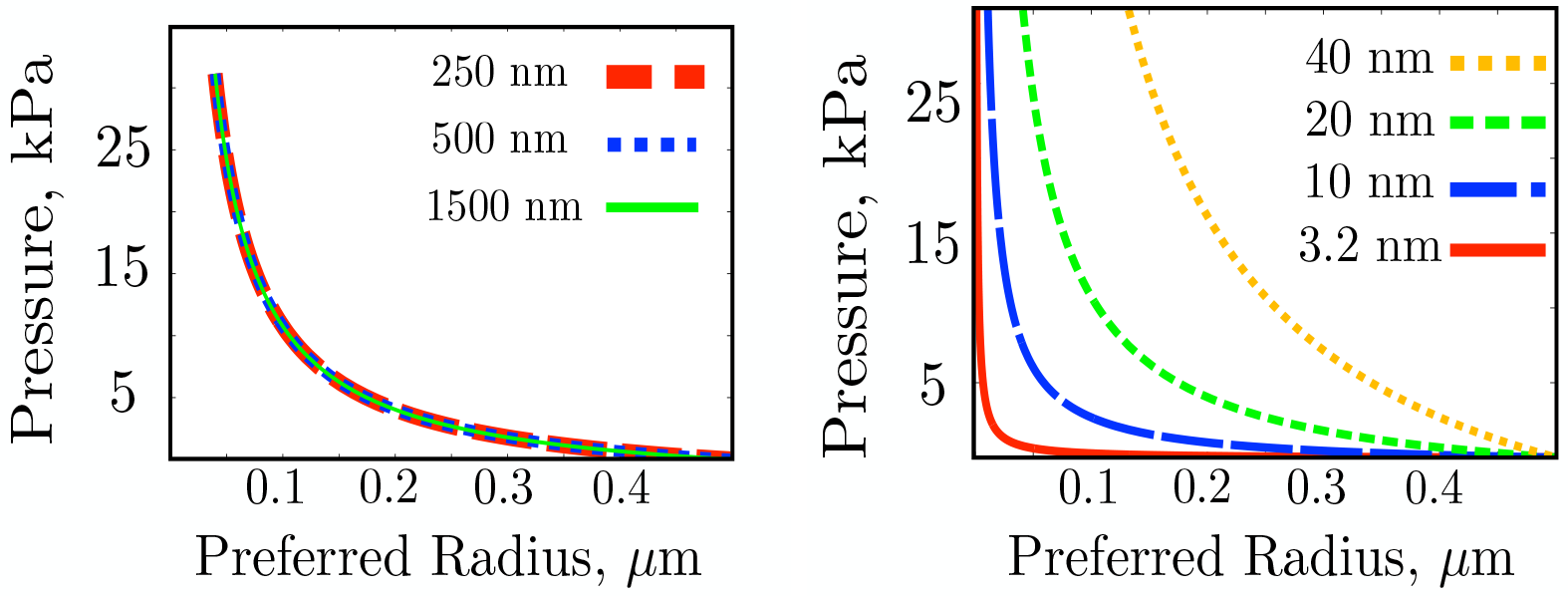
Pressure exerted vs preferred radius, for varying lengths as given in the inset, with bundle radius fixed at 20 nm (left). The length independence of the pressure exerted can be observed here. The pressure exerted against the preferred radius, with fixed bundle length taken 250 nm for different values of bundle radius as in the inset (right).

Given the effective internal pressure *P*_*eff*_, the effective stress in the hoop direction is *P*_*eff*_ *R*_*cell*_/*h*, where *R*_*cell*_ is the radius of the cell and *h* is the cell wall thickness. We explore the variation of the pressure exerted as the radius of the cell varies. Fixing *h*, we plot the pressure exerted against the radius of the cell (Figure 6 left). We observe that as *R*_*cell*_ increases, the inward pressure exerted by MreB becomes negligible. Interestingly, for *r* = 0.3*μm*, the pressure exerted attains maxima, when *R*_*cell*_ ≈ 0.5*μ*m. In general though, cell width is determined by a combination of multiple factors, for instance a delicate balance between the Rod system and class A penicillin binding proteins (aPBPs) [61]. In the following, we fix the preferred radius *r* of MreB to 0.3*μm*, noting that our results hold qualitatively over a wide range of values of *r* (0.2*μm* − 0.4*μm*).

**Figure 6:**
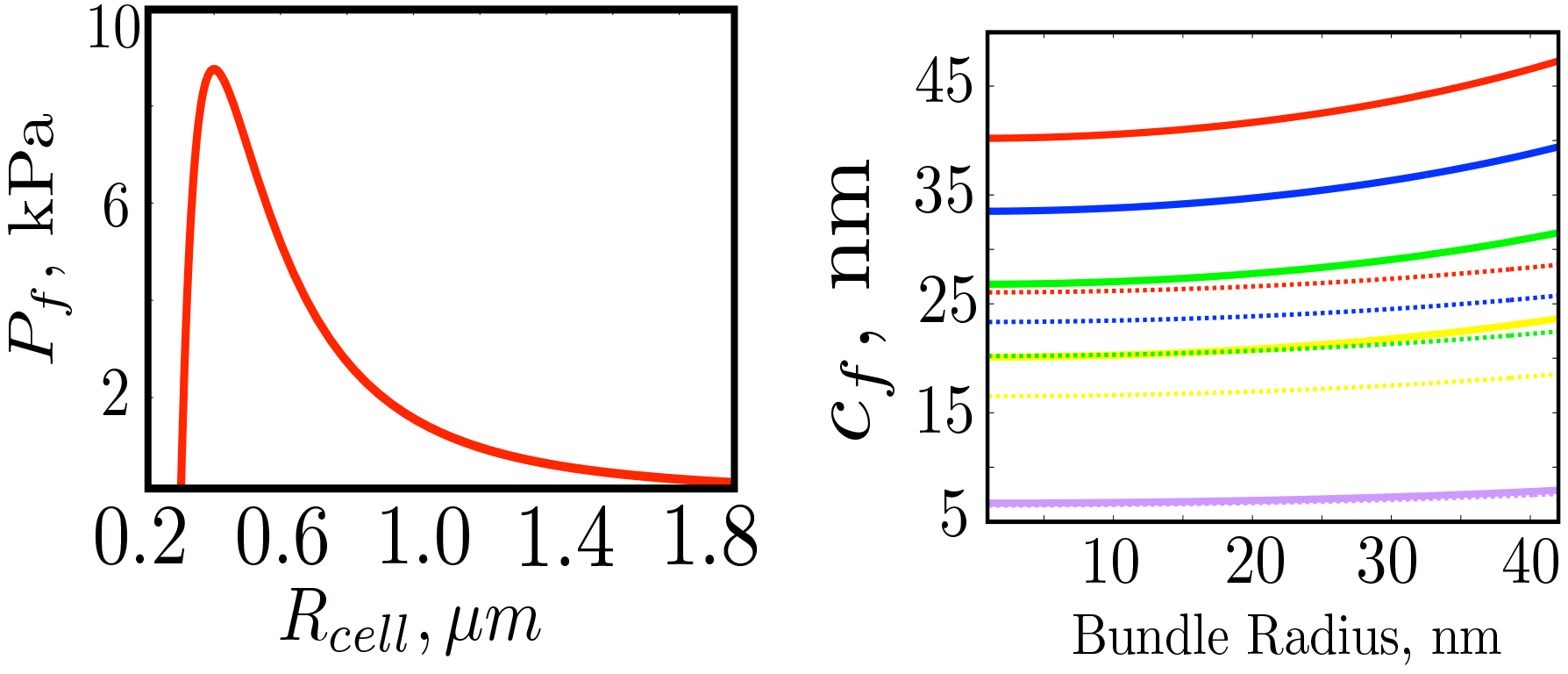
(Left) Average inward pressure exerted by MreB (right) with preferred radius *r* = 0.3*μ*m respectively. (Right) Critical crack length vs Bundle radius for MreB. Lines indicate the planar case, while dotted lines indicate the cylindrical case. In both cases, the degree of effect of crosslinking is taken as indicated, from bottom to top, n = 1 and n = 3, 4, 5, 6, depending on the length of the glycan strand.

### 4.2 Effect on crack length

We now quantify the role of MreB as a determinant of the critical crack length and compare its effect to that of the cross-linked structure of the cell wall. As explained previously, pressure is exerted by MreB in the inward direction. This acts to counter the turgor pressure, resulting in a lowering of the effective pressure on the cell wall. We now examine the effect of this lowering of the pressure, on the critical crack length of the cell wall. We plot the critical crack length against the bundle radius (Figure 6 (right)). We take into account the planar case and the cylindrical case, besides incorporating the effect of the cross-linking (with *n* as in Section 3.1). We note that when the bundle radius is zero, the effect of MreB is negated. So, the critical crack length in this case, is the same as the critical crack length as calculated in the first part, where the effect of MreB was not under consideraton. We observe that in all degrees of cross-linking, the critical crack length does not change much even for very large values of bundle radius. This underlines the significance of cross-linking in the protection of the cell wall against cracking.

It is however possible that MreB affects the toughness of the cell wall in other ways. For instance, a higher concentration of MreB might increase the cross-linking density [12], which can be deleterious for the toughness of the cell wall even as the stiffness of the cell will increase.

## 5 Discussion And Conclusions

In this work, we studied the role of the cross-linked structure of the cell wall in ensuring sufficient resistance to crack propagation. We deduced that the tearing energy varies inversely with the degree of cross-linking. We also showed that terminally cross-linked short length glycan strands can dramatically enhance the tearing energy. In particular, we showed that for about 30% cross-linking of the cell wall, as has been observed for *E.coli* [26], the optimal length of the glycan strands for maximizing the tearing energy, are shorter length glycan strands with length ∼ 7 - 8 disaccharides, cross-linked at the ends. This provides a possible explanation for surprising experimental observations, which have demonstrated an abundance of shorter length filaments in the peptidoglycan mesh [25] and of the strong preference of glycan strands to cross-link to each other at the termini [26, 27]. We used Griffith theory to calculate critical crack length for different degrees of cross-linking. Finally, we investigated the effect of MreB reinforcement of the cell wall, modelling MreB as several disconnected bent cylinders and estimated the inward pressure exerted for a wide range of parameters. We concluded that the effect of the cross-linked structure of the cell wall plays the primary role in ensuring the integrity of the cell wall.

An interesting feature of our analysis is the illustration of a standard dilemma faced when engineering materials, which is to to ensure optimal levels of stiffness and toughness, as the two requirements usually are at cross purposes [8] - in this case, we showed lower degrees of cross-linking result in higher tearing energies, thus offering better protection to the cell wall. On the other hand, a higher degree of cross-linking results in stiffer cell walls [12, 14], allowing the cell wall to bear turgor pressure and to preserve its shape. Similarly, longer glycan strands results in enhanced stiffness [29], while, shorter length glycan strands can amplify the toughness of the cell wall, as we exhibited. A natural question now is to understand how bacteria maintain an optimal degree of cross-linking, appropriate glycan strand length distribution and precise placing of the cross-links along the strand lengths, fine tuning their structure to ensure the right mix of mechanical properties under a variety of conditions. It will be particularly interesting here to probe the role of hydrolysis, which can affect both the degree of cross-linking and the glycan strand length distribution [15] and which, as we discussed, can mitigate the danger of failure due to crack propagation by cleaving appropriate peptide bridges.

The overarching theme of our analysis was to understand the molecular mechanistic underpinnings of the mechanical properties of the cell wall, in particular its toughness. More experiments are necessary to get a complete understanding of the problem, for instance analogous to those carried out for other biocomposites like nacre, which has an interesting sawtooth shaped force extension curve, explaining its remarkable toughness [62]. Further, experimental study of the behaviour of cracks on the cell wall under varying conditions can elucidate not just the mechanical properties of the cell, but also its growth process, as demostrated in recent experiments using laser nano-ablation, where cuts were introduced on *C.elegans* cell surfaces and were probed to study embryo elongation [34]. An outstanding question in this regard is to probe the ability of the bacterial cell to heal cracks on its surface, where invariably large scale remodelling of the cell wall has to be taken into account. A plausible method is that initially, perhaps counter-intuitively, peptide cross-links in the vicinity of the crack tip are cleaved by hydrolysis, which as our model explained, can act to locally hike the tearing energy and arrest the progress of the crack. Then wall material is inserted to repair the crack or to apply a mechanical force to close the crack. Since MreB directs peptidoglycan insertion [47], it naturally will play an important role in the crack repair process. In fact, it has been proposed that the observed MreB patch propagation in the circumferential direction is due to stable circumferential propagation of small gaps in the anisotropic sacculus [63]. Our model paves the way for more detailed work, leading upto a precise study of the mechanism by which cracks on the cell surface are healed even as the cell is growing. Such a mechanism will be fundamental to the survival of the cell and an understanding of this will require a comprehensive blend of experimental, computational and theoretical techniques, which can then be leveraged to design new age anti-bacterials.

## A Griffith Energy Balance Criterion

The foundation stone for fracture mechanics was laid by the work of Griffith, who studied the energetics of crack propagation [32]. When an increment in the length of the crack occurs, there is a release of strain energy. Griffith proposed that crack propagation becomes energetically favourable when this release of strain energy, due to increment in *crack* length, is greater than the surface energy which is needed to break the bonds of the specimen. The critical point is reached when both are balanced, which we write mathematically as

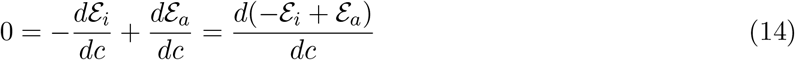

Here *ε*_*i*_ is the strain energy released, *ε*_*a*_ is the surface energy, *dc* denotes the crack length increment and we have that the total energy

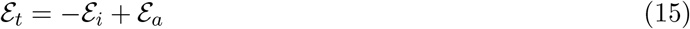

So, the critical crack length is the length at which the total energy attains its maxima. It is assumed that the thickness of the material is constant and that the crack growth is slow (quasi-static case). Further, it is assumed that energy dissipation, due to friction and plastic deformations, is negligible. The energy balance analysis of Griffith laid the foundation stone for linear elastic fracture mechanics (LEFM) and was key to the development of Irwin’s notion of stress intensity factors [64, 65]. In LEFM, energy dissipation, which occurs in a region surrounding the crack tip, called the dissipation zone or the process zone, is assumed negligible. But in many cases, the dissipative zone is quite large in size and LEFM proves inadequate, for which newer theories like Elastic Plastic Fracture Mechanics are used, including methods like J-integrals and cohesive zone model. For more details, we refer to [66, 38].

Let us now consider the case of a centrally placed crack of length 2*c* in an infinite plane of uniform thickness *h*, with a load *σ* applied in the direction perpendicular to the plane of propagation of the crack. The strain energy is given by *W* = ∫(*σ*: *ϵ*) *dV* where, *σ* is the stress tensor and *c* is the strain tensor. Assuming Hookean material with elastic modulus *E*, the strain energy density *E*, in the plane stress case, can be written as 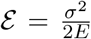 When the crack grows to length 2*c*, the strain energy released is then given by

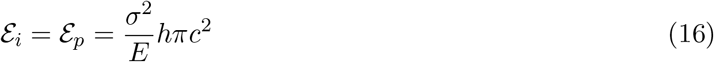

where *h* denotes the thickness of the specimen. Now, for the crack to propagate, the bonds in the plate have to be broken and hence, an amount of work equal to bond energy must be performed. Let *G*_0_ represent the minimum tearing energy (J/*m*^2^), required for the crack to propagate by breaking bonds in its path. Then, for a crack of length 2*c*, we have

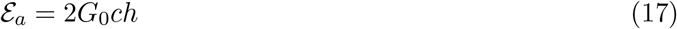

Substituting *ε*_*p*_ and *ε*_*a*_ in Eq 14 appropriately, we get

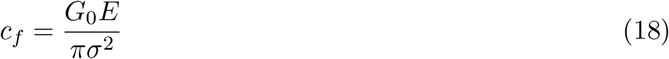

is the Griffith critical crack length, which is interpreted as the length above which a crack will grow uncontrollably, when the loading at the edges is fixed at *σ*.

## B Crack on orthotropic cell wall

Here, we calculate the strain energy released for a crack in orthotropic model of the cell wall and compare it with the isotropic case.

For plane stress in the orthotropic case, by Hooke’s law, we have

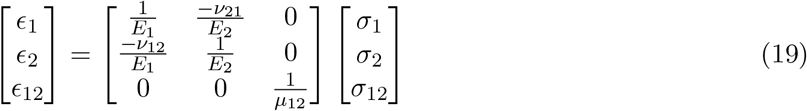

where *E*_1_, *E*_2_ denote the elastic modulus, *ν*_12_, *ν*_21_ denote the Poisson’s ratio and *μ*_12_ denotes the shear modulus.

We consider an infinite orthotropic plate of thickness *h* placed in the *Y Z*-plane, with a crack of length 2*c* placed in the plate, centred at the origin and along the *Z*-axis, as in Figure 2(a). Fixing *Z* = 1 and *Y* = 2, we get the strain energy released as [67]

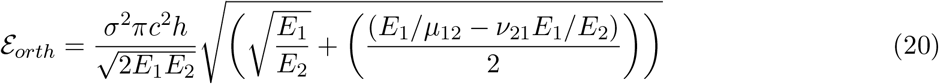

which we rewrite as (cf. 3)

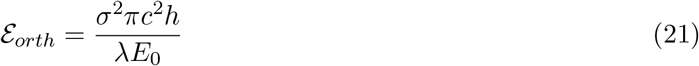

where 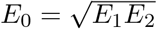 and 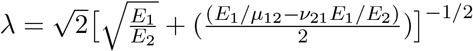

In our case, we have *E*_1_ ≈ 25 MPa, *E*_2_ ≈ 45 MPa, *ν*_21_ ≈ 0.46, *ν*_12_ ≈ 0.2 [23]. However, the shear modulus *μ*_12_ of the cell wall is not clear. So, we assume that the cell wall is “specially” orthotropic, which gives 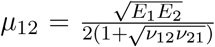. In this case, we have *λE*_0_ ≈ 36 MPa, comparable to the value of *E* = 30 MPa that is used to model the cell wall as an isotropic material implying that the strain energy released are quantitatively similar in both cases. So, our model framework and the main conclusion of our paper on the role of the cross-linked structure of peptidoglycan mesh in increasing the tearing energy of the cell wall does not change with the inclusion of anisotropy in the model.

## C Curvature effect on Strain energy

We now calculate the effect of geometry on the strain energy released due to a through the thickness crack of length 2*c*, aligned longitudinally on a pressurized thin cylindrical shell. In this case, apart from in-plane deformations, there is an additional out-of-plane deformation in a small region around the crack resulting from the action of normal force due to internal pressure which is directed through the shell.

The deflection *δ* in the normal direction, which in this case is the radial direction, varies over the distance *l*. The bending energy per unit area is given by 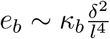, where we have used that the curvature change is of the order 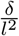 [58]. The total bending contribution (ignoring all coefficients) concentrated over the area *l*^2^ is then given by

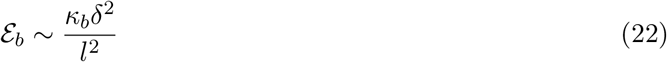

where, *κ*_*b*_ ≈ *Eh*^3^ is the bending rigidity.

The strain tensor for the stretching energy is of the order of 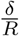, that is 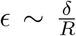, where *R* is the radius of the cylinder. The corresponding stress is then 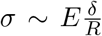 and the stretching energy per unit area is given by 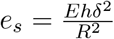. Thus, the total stretching energy contribution is (ignoring all coefficients)

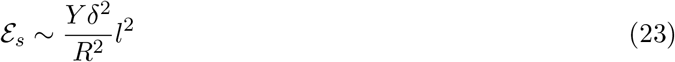

where, *Y* = *Eh* is the 2-*D* Young’s modulus for the case of plane stress.

The curvature correction energy term *ε*_*cyl*_ is therefore given by

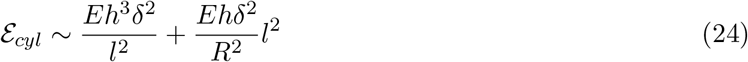

Minimizing the total energy in Equation 24, 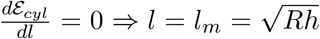 which gives us a new elastic length scale for localization of deformation.

Plugging in the value of *l*_*m*_ obtained above, we get

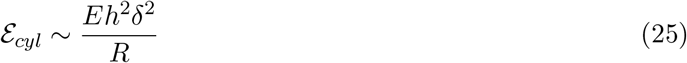

We denote the force acting on the crack periphery, due to internal pressure, by *f*. Varying *ε*_*cyl*_ with respect to *δ* and equating it to the work done by the force *f*, we get

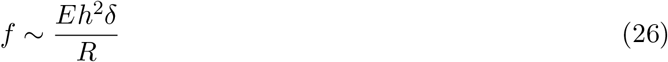

The force acts along the line of the crack, and hence, the area over which normal stress is applied is of the order of *c*^2^, where 2*c* is the crack length. It follows then that the force acting is of the order of *f*∼ *Pc*^2^, where *P* is the internal pressure. Substituting this value of *f* in equation 26, we get the normal deflection 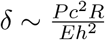. Therefore, using Equation-25, we get

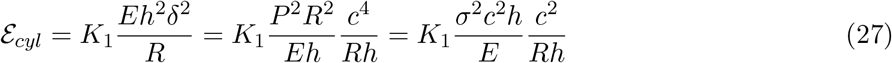

where *K*_1_ is a dimensionless constant. This constant 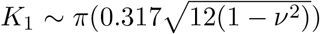 [35, 68], where *ν* denotes the Poisson’s ratio of the shell material. Since 0 ≤ *ν* ≤ 1/2 [13, 23], we have that *K*_1_ ≈ *π*. Therefore, we get that the total strain energy released is given by

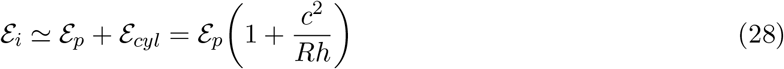

## D Glycan Strand length and Tearing energy

In this section, we relate the glycan strand lengths to the tearing energy of the cell wall. Our objective here is to estimate *n*, the average number of glycosidic bonds connecting disaccharides units between adjacent cross-links, as defined in Section 3.1, which we relate here to the glycan strand length. For any glycan strand denoted *g*, this value is given by

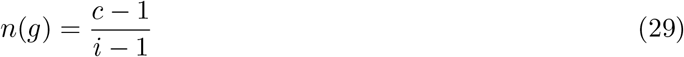

where *c*(*g*) denotes the total number of disaccharide units between the two extreme cross-links of *g* (in other words, we are counting those disaccharide units in the glycan strand that lie between any pair of adjacent cross-links) and *i*(*g*) denotes the total number of cross-linked peptide stems on the glycan strand *g*.

We first consider the case when glycan strand lengths span the circumference of the cell wall. In this case, due to periodicity, *c*(*g*) = *l*(*g*), where *l*(*g*) denotes the total number of disaccharide units for glycan strand *g*. Since *l* is large, it follows that *n* ∼ 1/*k* where *k* denotes the fraction of cross-linked peptides. In case of *E.coli*, *k* ∼ 0.3 [69], so then *n* ∼ 3. This also illustrates how a lower degree of cross-linking can enhance the toughness of the cell wall.

However, we now show how smaller length glycan strands cross-linked at the ends, can considerably increase value of *n*. Let us consider a glycan strand *g* of length *l* disaccharide units. It follows from Equation 29 that the value of *n* for a strand of fixed length will be maximized when the peptide stems at the end of the glycan strands are cross-linked. In this case, *c*(*g*) ∼ *l*. Also, for the degree of cross-linking denoted *k*, we will have *i* ∼ *kl*. Therefore,

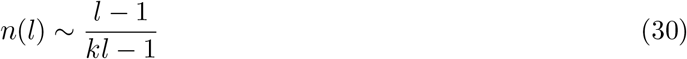

Taking *k* = 0:3 and allowing for atleast two cross-links across the length of the strand, we get *n* ∼ 3 - 6. The value of *n* decreases as the length of the glycan strand increases and for length of glycan strand *l*(*g*) ∼ 7 - 8, the value of *n* ∼ 5 - 6. Now, given a glycan strand length distribution *p*, the average value of *n* across the cell wall is

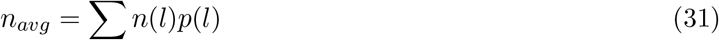

where *p*(*l*) denotes the proportion of glycan strand lengths of length *l*. It is clear that a higher proportion of glycan strands of strand lengths ∼ 7 - 8 disaccharide units will enhance the average value of *n* across the cell wall. For very short strands with length 4 disaccharides or less, only a single peptide stem will be cross-linked with cross-linking degree around 30% [69], so a similar analysis is not possible. It is not clear what role these very short strands play in the peptidoglycan mesh.

## E Dissipative Zone

The length of the dissipative zone, denoted by *l*_*d*_, is given by

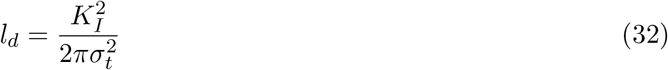

where *K*_*I*_ is the stress intensity factor and *σ*_*t*_ is the tensile strength, which is the maximum tensile stress a body can take before failure [44, 9, 38]. We have the strain energy release rate 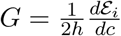 and the stress intensity factor is related to the strain energy release rate as [38]

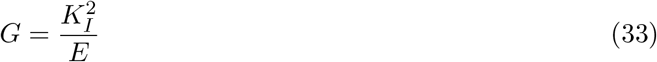

Now, denoting by ∆*y*_*t*_ the maximum stretching of glycan strands, it follows from force balance that

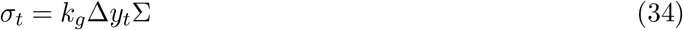

where *k*_*g*_ denotes the spring constant of the glycan strands, which is given by

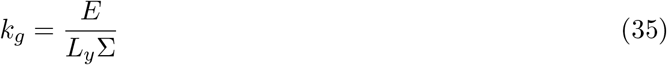

where *L*_*y*_ denotes the length of the glycan strands between adjacent crosslinks. Also, minimum tearing energy, *G*_0_ can be written in terms of *k*_*g*_ as

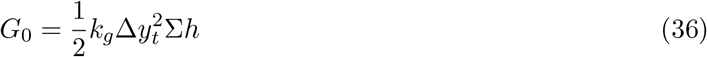

So, it now follows from Equations 32, 33, 34, 35, 36 that

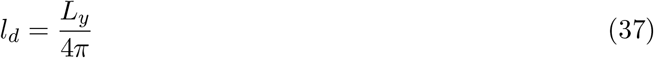

Since the typical length of the glycan strand between adjacent cross-links is ∼ 3-6 disaccharides, we have that *l*_*d*_ ≈ 5Å.

## F Curves on Surfaces and the Darboux Frame

Let 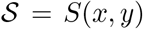 be a surface in ℝ^3^. We denote the tangent vectors *S*_*x*_ and *S*_*y*_, which span the tangent plane to the surface at the *S*(*x, y*). Suppose now that we have a curve on the surface, given by the mapping

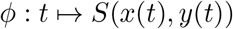

The tangent to the curve at the point *S*(*x*(*t*), *y*(*t*)) is then given by 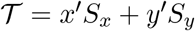. We now define a frame for the curve *ϕ* (which is thought of as the centerline of the filament bundles under consideration in this work). In this case, we want to use frame to incorporate the information that the curve lies on a surface, thus we will be using the Darboux frame and not the usual Frenet-Serret frame.

Let 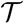 denote the tangent to the curve at *S*(*x*(*t*), *y*(*t*)) as before. Let 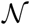 denote the normal to the surface at *S*(*x*(*t*), *y*(*t*)). We also further define another unit vector 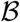 as 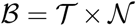. Together, the vectors 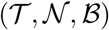 define the darboux frame for the curve *ϕ*, lying on the surface 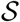.

It is easy to see, since 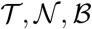 are orthonormal, we have that 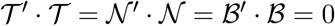 and further

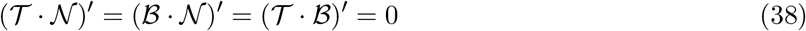

**Figure 7:**
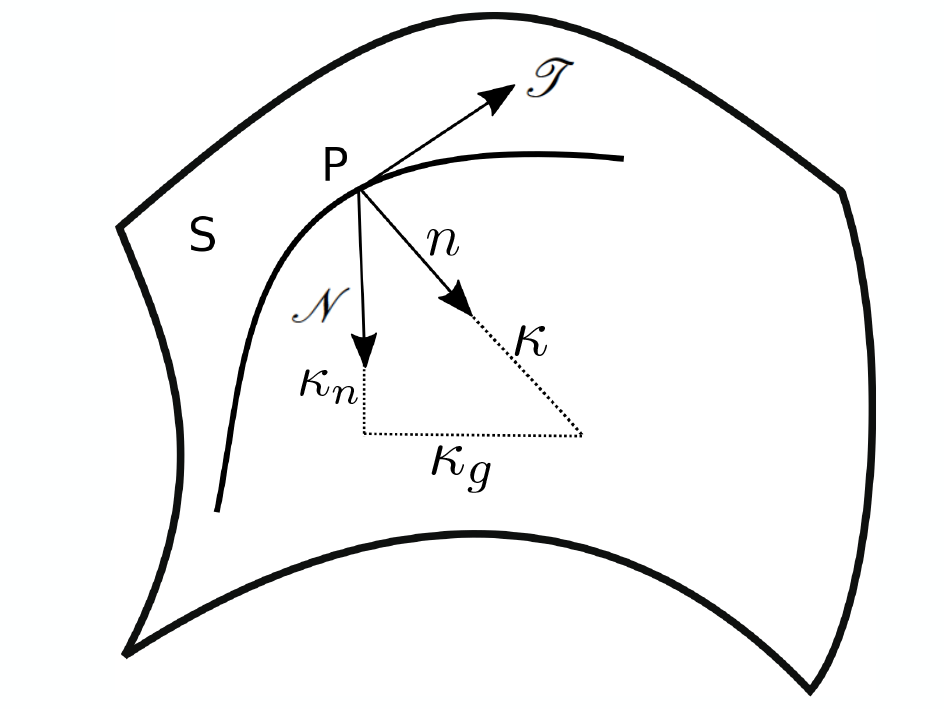
The Darboux frame describing curves on surfaces, with 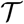 being the tangent to the curve, 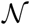 the surface normal. The Darboux frame is 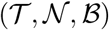, where 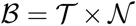.

This implies the existence of scalar functions *κ*_*n*_, *κ*_*g*_ and *τ* such that we have

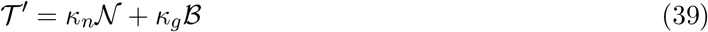

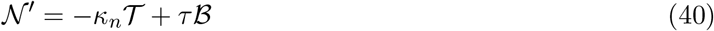

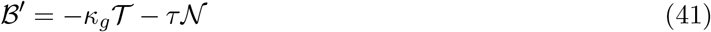

Here, *κ*_*n*_ is the normal curvature of the curve, *κ*_*g*_ is the geodesic curvature of the curve and *τ* is the twist of the curve.

We can now define the darboux vector

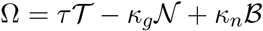

and thus, we will have

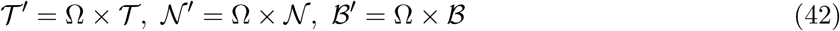

## G MreB filament bundles in the cell

We estimate the number of MreB filament bundles in a typical cell. The number of MreB monomers has been estimated to lie in the range 17000 - 40000 (see Table 2). The number of filament bundles will depend on the number of MreB monomers in a single bundle, which in turn will depend on the radius of the bundle, its length and the packing of the monomers in the bundle. Since the bundle is modelled as a cylinder of length *L*_*fil*_ = *l*_*f*_ and radius *a*, and monomers are modelled as balls of fixed radius *r*_0_ (see Fig. 8), the number of monomers in the filament bundle will be a fraction of number 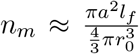. This is effectively a problem of packing of balls of uniform radius in a cylinder of given dimensions. It is obvious that not all the volume of the cylinder can be occupied by the balls. Following the resolution of Kepler’s conjecture, it is now known that for sufficiently large containers, the volume fraction occupied by balls of a uniform small radius is bounded above by 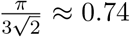, which is achieved by cubic close packing or hexagonal close packing [59]. In other words, the densest packings occupy about 0.74 of the volume of the container.

**Figure 8:**
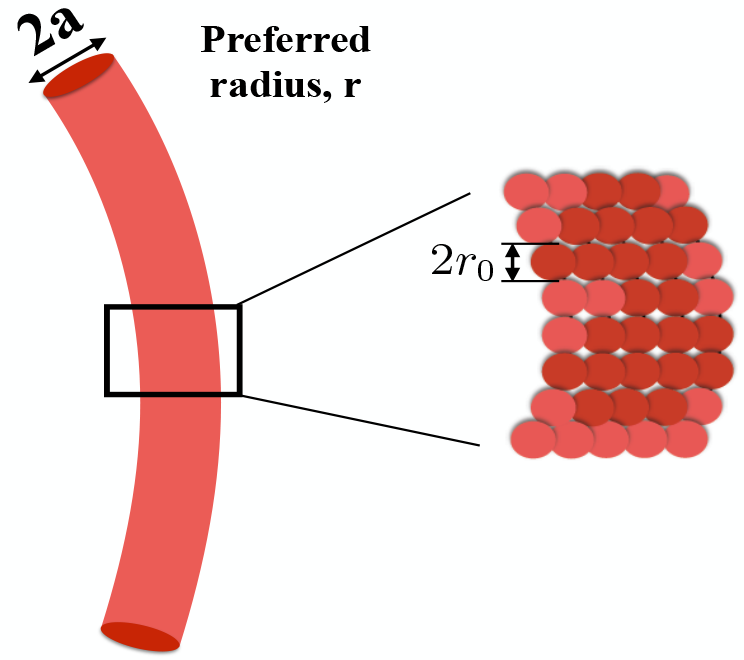
A MreB filament bundle with bundle radius *a* and preferred radius *r*. The filament bundle consists of monomers modelled as spheres of radius *r*_0_.

Another possible configuration can be described as follows- we can think of each filament bundle as an aggregation of filaments with each filament having the same number of monomers *∼ l*_*f*_ /2*r*_0_. Calculating the number of filaments will thus give us the total number of monomers in this configuration. This is given approximately by dividing the area of cross-sectional circle of the cylindrical container by the area of the equatorial circle of the monomer, giving us the number 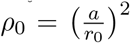. Here we are assuming that the arrangement is such that the equatorial circles of the monomers on the top cover almost the whole area of the cross-sectional circle of the cylinder and that the monomers are arranged in collection of straight lines, piled below the monomers on the top. Thus, the number of monomers in this arrangement is given by 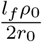 and it is straightforward to see that in this case, the monomer balls occupy ∼ 2/3 of the volume of the cylinder.

Now, since the filaments have a robust elastic modulus of around 2 GPa, it is fair to assume to that the packing of the monomers has to be sufficiently dense. On the other hand, since the monomers are much smaller than the cylindrical filament, the upper bound for the volume has to be around 0.74, as explained above. So, we assume that the volume fraction of the monomers in the cylindrical container is ∼ 0.5 - 0.75. So, with bundle radius denoted *a* and length denoted *l*_*f*_, and with total number of MreB monomers denoted *N*, we have that the number of filaments *n*_*f*_ lies in the range 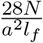 to 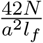 with a and *l*_*f*_ given in nanometers and where we have used the value *r*_0_ = 2.5 nm (Table 2). We tabulate (Table 1) the number of filament bundles, where the length of such filament bundles taken as 250 nm, 500 nm and 1500 nm, and bundle radius taken as 3.2 nm, 10 nm, 20 nm and 40 nm. We have assumed here, for simplicity, that all the MreB monomers in the cell are part of some MreB bundle. However, there can be several MreB monomers in the cell cytoplasm, so that the values in Table 1 give us an upper bound for the number of MreB bundles present in the cell.

## H Helix model of MreB

We now specialize to the case where MreB assumes a cell spanning helical configuration. We consider a circular cylinder of radius *R*. For a helix of pitch, *p* = 2*πB* constrained to this cylinder, we have *κ*_*g*_ = 0, *κ*_*n*_ = *R/*(*R*^2^ + *B*^2^) and *τ* = *B/*(*R*^2^ + *B*^2^). We assume that the preferred configuration of MreB is helical, with curvatures denoted *c*_*n*_ = *r/*(*r*^2^ + *b*^2^) and *c*_*t*_ = *b/*(*r*^2^ + *b*^2^), where *r* and *p*_0_ = 2*πb* are the preferred radius and pitch respectively. It follows now from Eqn. 9 that

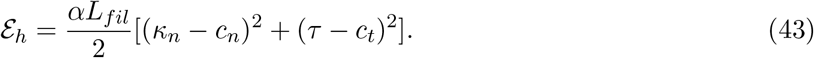

To simplify calculations, we introduce the helix opening angle *θ*, defined as the fixed angle for which, if *s* is the arc length of the helix, then *z* = *s* sin *θ*, where *z* is the *Z*-axis coordinate. We will assume that the helix is left-handed [70]. Thus, 0 *≤ θ ≤ π/2*. Further,

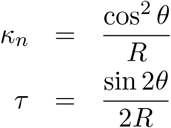

with *R* denoting the radius of the cylinder. We also have *L*_*fil*_ = *L/* sin *θ* where *L* is the length of the cylinder.

To find the final configuration of the helix on the cylinder, we now differentiate *E*_*h*_ with respect to *θ*, then equate it to zero, to get the following equation

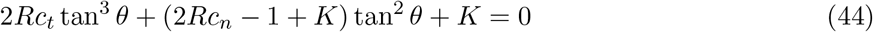

where

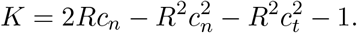

This equation has a unique positive solution, which gives the opening angle *θ*_0_, yielding the final pitch.

More generally, the roots of this equation can be used to derive a relation between the final pitch and the preferred pitch, for varying preferred radius. We plot this in Fig. 9 (Left), where the final pitch is mapped against the preferred pitch. Fluorescence imaging of MreB suggests that the helix makes around 1-2 turns per *μ*m [71, 72], which gives the pitch of the helix to be in the range 0.5*μ*m-1*μ*m. We then have, from Fig. 9 (Left), that the preferred radius must be ∼ 0.4*μm*. We take the pitch of the helix to be *≈* 0.8*μ*m. The preferred pitch of the helix is then *≈* 0.5*μ*m (seeFigure 9 (Left))

The average pressure exerted on the cell wall can now be calculated as in Section 4.1, which is given by

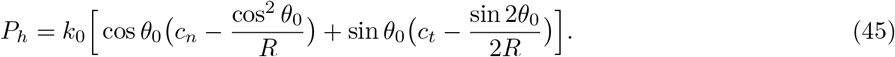

where 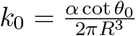. Note that *P*_*h*_ ∝ *a*^4^, where *a* denotes the bundle radius, so the pressure exerted increases sharply as the bundle radius increases. It can be shown that for a wide range of the helix bundle radius (5*nm* − 40*nm*) [12, 71], the pressure exerted shows similar low values of pressure exerted as in the filament bundle model.

**Figure 9:**
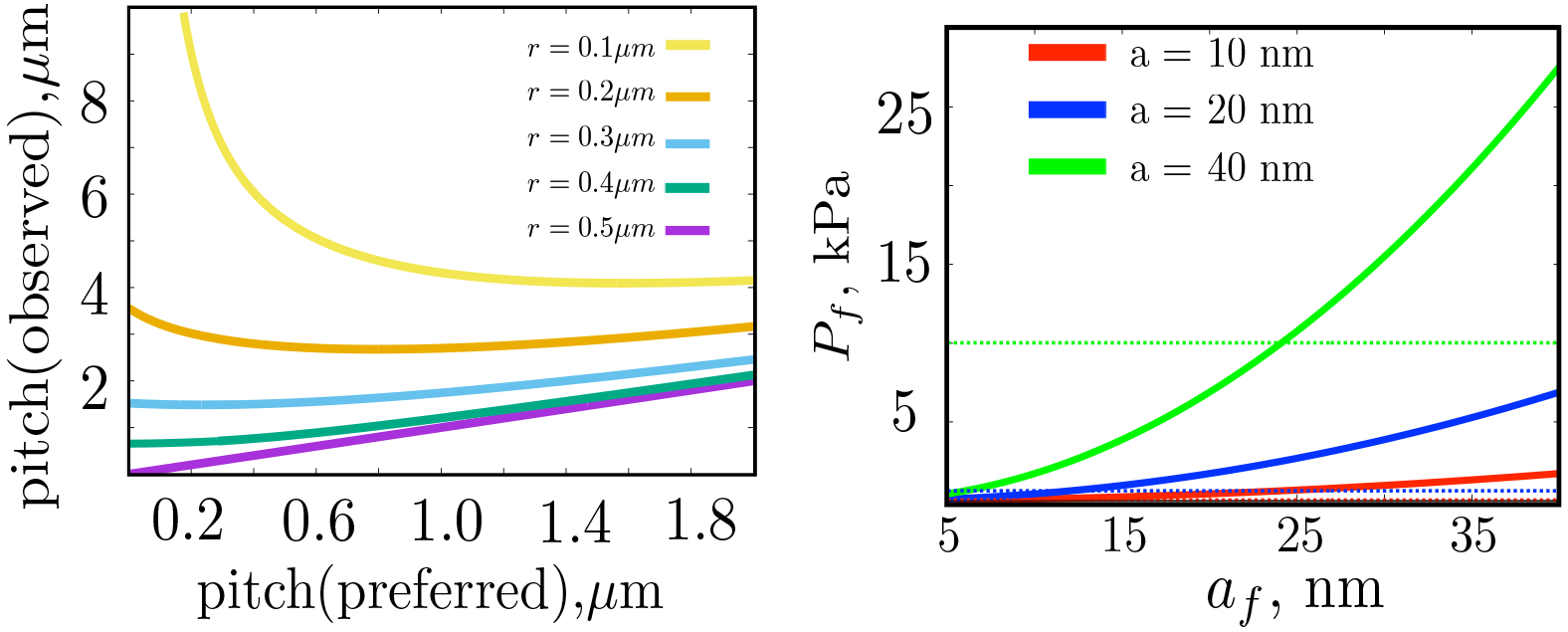
(left) The pitch, obtained from solving Eqn. 44, against the preferred pitch, *p*_0_ for varying preferred radius. It is observed that only for the higher values of the preferred radius, viz. ≥ 0.4*μm*, is the observed pitch *p* in the range 0.5*μ*m-1*μ*m. (right) A comparison of the inward pressure exerted in the helix model and the filament bundle model. As explained in the text, we calculate the number of MreB monomers in cell spanning helix with bundle radius, from bottom to top, 10 nm, 20 nm, and 40 nm. This is then used to calculate the number of filament bundles, the radius of which is picked from the x-axis and the length is fixed at 250 nm. The inward pressure exerted in this case is then calculated, it is the corresponding y-coordinate and is plotted as solid lines. The straight dotted lines indicate the pressure exerted by the helix of labelled bundle radius.

A natural question is to compare the inward directed pressure for the two models of MreB. An appropriate comparison can be made if we consider the two cases when the number of MreB molecules are the same, as are the packing conditions. So, we first estimate the number of MreB monomers (say *N_h_*) in the helical structure of appropriate radius, under same packing conditions as in the case of filament bundle. We then calculate the number of MreB bundles of fixed length and given radius under the assumption that the cell has *N*_*h*_ MreB monomers, which is then used to calculate the pressure exerted. More specifically, given a helix of pitch *p* and radius *a* = *a*_*h*_, its volume is given by

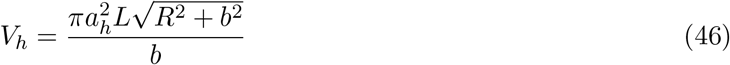

where *L* denotes the length of the cell, *R* denotes the radius of the cell and *b* = *p/*2*π*. Under the assumption that the volume occupied by the balls representing the MreB monomers (each of radius *r*_0_ = 2.5 nm), is half the volume of helical bundle, we have *N*_*h*_ ≈ *V*_*h*_/2*V*_*MreB*_, where 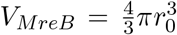. We then calculate the number of bundles, denoted *n*_*f*_, of radius *a*_*f*_ and length *l*_*f*_, under the assumption that there are *N*_*h*_ MreB monomers in the cell, again assuming that the volume occupied by MreB monomer balls is half of the volume of the filament bundles. As explained in Appendix G, we will then have 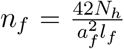. This is now used to calculate the pressure exerted inwards, and to compare with the helix case, as in Figure 9 (right), where we plot pressure exerted against the MreB bundle radius, with bundles of fixed length and the number of such bundles calculated as above. The colors in Figure 9 (right) indicate the helix bundle radius, while the straight dotted lines indicate the pressure exerted by the helix of given bundle radius. It is clear that in each case, the pressure exerted by the filament bundles is more than the helix. To illustrate this, let us consider the case of a helical bundle of radius 40 nm (represented by color green in Figure 9 (right)). We can observe that, in this case, even for relatively low values of the filament bundle radius, of around 20 nm, the pressure exerted is already matching the pressure exerted by the helix, while at around a bundle radius of 40 nm, the pressure exerted far exceeds the pressure exerted by the helix. It therefore follows from Figure 9 (right) that having several shorter length filament bundles exerts higher inward directed pressure than a helical bundle under similar conditions.

